# ideal: an R/Bioconductor package for Interactive Differential Expression Analysis

**DOI:** 10.1101/2020.01.10.901652

**Authors:** Federico Marini, Jan Linke, Harald Binder

## Abstract

**Background:** RNA sequencing (RNA-seq) is an ever increasingly popular tool for transcriptome profiling. A key point to make the best use of the available data is to provide software tools that are easy to use but still provide flexibility and transparency in the adopted methods. Despite the availability of many packages focused on detecting differential expression, a method to streamline this type of bioinformatics analysis in a comprehensive, accessible, and reproducible way is lacking.

**Results:** We developed the `ideal` software package, which serves as a web application for interactive and reproducible RNA-seq analysis, while producing a wealth of visualizations to facilitate data interpretation. ideal is implemented in R using the Shiny framework, and is fully integrated with the existing core structures of the Bioconductor project. Users can perform the essential steps of the differential expression analysis workflow in an assisted way, and generate a broad spectrum of publication-ready outputs, including diagnostic and summary visualizations in each module, all the way down to functional analysis. ideal also offers the possibility to seamlessly generate a full HTML report for storing and sharing results together with code for reproducibility.

**Conclusion:** ideal is distributed as an R package in the Bioconductor project (http://bioconductor.org/packages/ideal/), and provides a solution for performing interactive and reproducible analyses of summarized RNA-seq expression data, empowering researchers with many different profiles (life scientists, clinicians, but also experienced bioinformaticians) to make the *ideal* use of the data at hand.

## Background

Over the last decade, RNA sequencing (RNA-seq, (Wang *et al.*, 2009)) has become the standard experimental approach for accurately profiling gene expression. Complex biological questions can be addressed, also thanks to the development of specialized software for data analysis; these aspects are, e.g., reviewed in the works of Conesa *et al.* (Conesa *et al.*, 2016) and Van den Berge *et al.* (Van den Berge *et al.*, 2019), which cover a broad spectrum of the possible applications.

Differential expression analysis is a very commonly used workflow (Oshlack *et al.*, 2010; Love *et al.*, 2015; Soneson *et al.*, 2015; Ritchie *et al.*, 2015), whereby researchers seek to define the mechanisms for transcriptional regulation, enabled by the comparisons between, for example, different conditions, genotypes, tissues, cell types, or time points. The ultimate aim is to determine robust sets of genes that display changes in expression, and to contextualize them at the level of molecular pathways, in a way that can explain the biological systems under investigation and provide actionable insights in basic research and clinical settings (Beigh, 2016).

Established end-to-end analysis procedures (such as (Anders *et al.*, 2013; Pertea *et al.*, 2016; Love *et al.*, 2018)) are nowadays available, yet often bioinformatics analyses can be a challenging and time-consuming bottleneck, especially for users whose programming skills do not suffice to flexibly combine and customize the steps (and software modules) of a complete analysis pipeline.

Current software implementations for quality assessment (e.g. FastQC, https://www.bioinformatics.babraham.ac.uk/projects/fastqc/), preprocessing, alignment (Dobin *et al.*, 2013), and quantification (Anders *et al.*, 2015; Liao *et al.*, 2014; Patro *et al.*, 2017; Bray *et al.*, 2016) have streamlined the generation of large matrices of the transcriptome profiles. These intermediate results have to be provided as input to software for differential expression analysis (Love *et al.*, 2014; McCarthy *et al.*, 2012; Ritchie *et al.*, 2015), which constitute core components of the R/Bioconductor project (Gentleman *et al.*, 2004; Huber *et al.*, 2015).

In our previous work (Poplawski *et al.*, 2016), we reviewed a selection of interfaces for RNA-seq analysis from the perspective of a life scientist, defining criteria that cover many essential aspects of every software/framework, including e.g. installation, usability, flexibility, hardware requirements, and reproducibility. Building upon these results, we subsequently developed a tool that satisfies a broad set of requirements for differential expression analysis and is presented in the following.

As a result of close collaborations with wet-lab life scientists and clinicians, we developed our proposal as an interactive Shiny (Chang *et al.*, 2016) web based application in the `ideal` R/Bioconductor package, which guides the user through all operations in a complete differential expression analysis. `ideal` provides an integrated platform for extracting, visualizing, interpreting, and sharing RNA-seq datasets, similar to what our Bioconductor `pcaExplorer` package does for the fundamental step of exploratory data analysis (Marini and Binder, 2019).

The `ideal` package takes as input a count matrix and the experimental design information, for allowing to also analyze complex designs (such as multifactorial experimental setups), while making it easy to reproduce and share the analysis sessions, promoting effective collaboration between scientists with different skill sets, an open research culture (Nosek *et al.*, 2015), and the adherence to the FAIR Guiding Principles for scientific data (Wilkinson *et al.*, 2016). Moreover, `ideal` delivers a wide range of information-rich visualizations, charts, and tables, both for diagnostic and downstream steps, which taken together form a comprehensive, transparent, and reproducible analysis of RNA-seq data.

`ideal` reprises and expands some design choices of `pcaExplorer`, with an improved documentation system based on tooltips and on the adoption of self-paced learning tours of the main functionality (with the `rintrojs` library (Ganz, 2016)). State saving and automated HTML report generation via knitr and R Markdown, following a template bundled in the package itself (which can be edited by the experienced users to address specific questions), ensure code reproducibility, which has received increasing attention in recent years (Peng, 2011; McNutt, 2014; Stodden *et al.*, 2016; Marini and Binder, 2016; Eglen *et al.*, 2017; Perkel, 2018).

A growing number of software packages (Younesy *et al.*, 2015; Nelson *et al.*, 2016; Su *et al.*, 2017; Harshbarger *et al.*, 2017; Gardeux *et al.*, 2017; Lim *et al.*, 2017; Li and Andrade, 2017; Zhu *et al.*, 2018; Ge *et al.*, 2018; Monier *et al.*, 2018; McDermaid *et al.*, 2018; Schultheis *et al.*, 2018; Kucukural *et al.*, 2019; Choi and Ratner, 2019; Price *et al.*, 2019; Tintori *et al.*, 2020; Su *et al.*, 2019) have been developed to operate on tabular-like summarized expression data, or on formats which might derive from their results (see Additional file 1: Table S1 for a comprehensive list of details on their functionality and characteristics).

Similar to `ideal`, many of the existing tools accept count data in tabular format, and proceed to compute differentially expressed genes, accompanying this with visualizations (both focused on samples and on features) and sometimes downstream operations such as functional analysis for the identified subset. In most of the existing software, single genomic features of interest can be inspected, with some support for identifier conversion. The majority of these solutions are distributed as standalone web applications (commonly in R/Shiny, although some are written in Javascript); still, not all of these can be easily distributed as packages, or deployed seamlessly to private local instances. While still underrepresented, some of the tools allow the generation of an analysis product, which in many cases is based on a report in R Markdown (Allaire *et al.*, 2018), dynamically generated at runtime.

Overall, the existence of many such software packages highlights the need for a user-friendly framework to generate rich outputs for assisting analysis and interpretation, yet currently none of the existing proposals is offering the complete set of features we implemented in our work, with a full integration in the Bioconductor environment, and a seamless combination of interactivity and reproducibility.

The `ideal` package integrates and connects a number of R/Bioconductor packages, wrapping the current best practices in RNA-seq data analysis with a coherent user interface, and can deliver multiple types of outputs and visualizations to easily translate transcriptomic datasets into knowledge and insights. By leveraging the efficient core structures of Bioconductor, `ideal` allows flexible additional visualizations, as it is possible with custom scripts or with other GUI-based tools such as the `iSEE` package (Rue-Albrecht *et al.*, 2018; Amezquita *et al.*, 2019).

`ideal` is available at http://bioconductor.org/packages/ideal/, and the application can additionally be deployed as a standalone web-service, as we did for the publicly hosted version available at http://shiny.imbei.uni-mainz.de:3838/ideal, where the readers can explore the functionality of the app.

## Implementation

### General design of ideal

`ideal` is written in the R programming language, wiring together the functionality of a number of widely used packages available from Bioconductor. `ideal` uses the framework of the `DESeq2` package to generate the results for the Differential Expression (DE) step, as it was found to be among the best performing in many experimental settings for simple and complex eukaryotes (Schurch *et al.*, 2016; Froussios *et al.*, 2019). Internally, this framework includes the estimation of size factors (with the median ratio method) and of the dispersion parameters, followed by the generalized linear model fitting and testing itself.

The web application and all its features are provided by a call to the `ideal()` function, which fully exploits the Shiny reactive programming paradigm to efficiently (re-)generate the rendered components and outputs upon detection of changes in the input widgets.

The layout of the user interface is built on the `shinydashboard` package (Chang and Borges Ribeiro, 2018), with a sidebar containing widgets for the general options, and the main panel structured in different tabs that mirror the different steps to undertake to perform a comprehensive differential expression analysis, from data setup to generating a full report. The task menu in the dashboard header contains buttons for state saving, either as binary RData files or as environments in the interactive workspace, accessible after closing the app.

Alongside features like tooltips, based on the bootstrap components in the `shinyBS` package (Bailey, 2015), `ideal` uses collapsible elements containing text to quickly introduce the functionality of the diverse modules, and guided tours of the user interface via the `rintrojs` package (Ganz, 2016), which provide means for learning-by-doing by inviting the user to perform actions that reflect typical use cases in each section. The *Quick viewer* widget in the sidebar keeps track of the essential objects, which are either provided upon launch, or computed at runtime, while `valueBox` elements (whose color turns from red to green once the corresponding object is available) above the main panel display a brief summary of each.

We invested particular attention in designing the application to guide the user through the different workflow steps (Figure 1). This can be appreciated in the steps for the *Data Setup* panel, which appear dynamically once the required input and parameters are provided. Moreover, we used conditional panels to activate the functionality of each tab only if the underlying objects are available.

**Figure 1:**
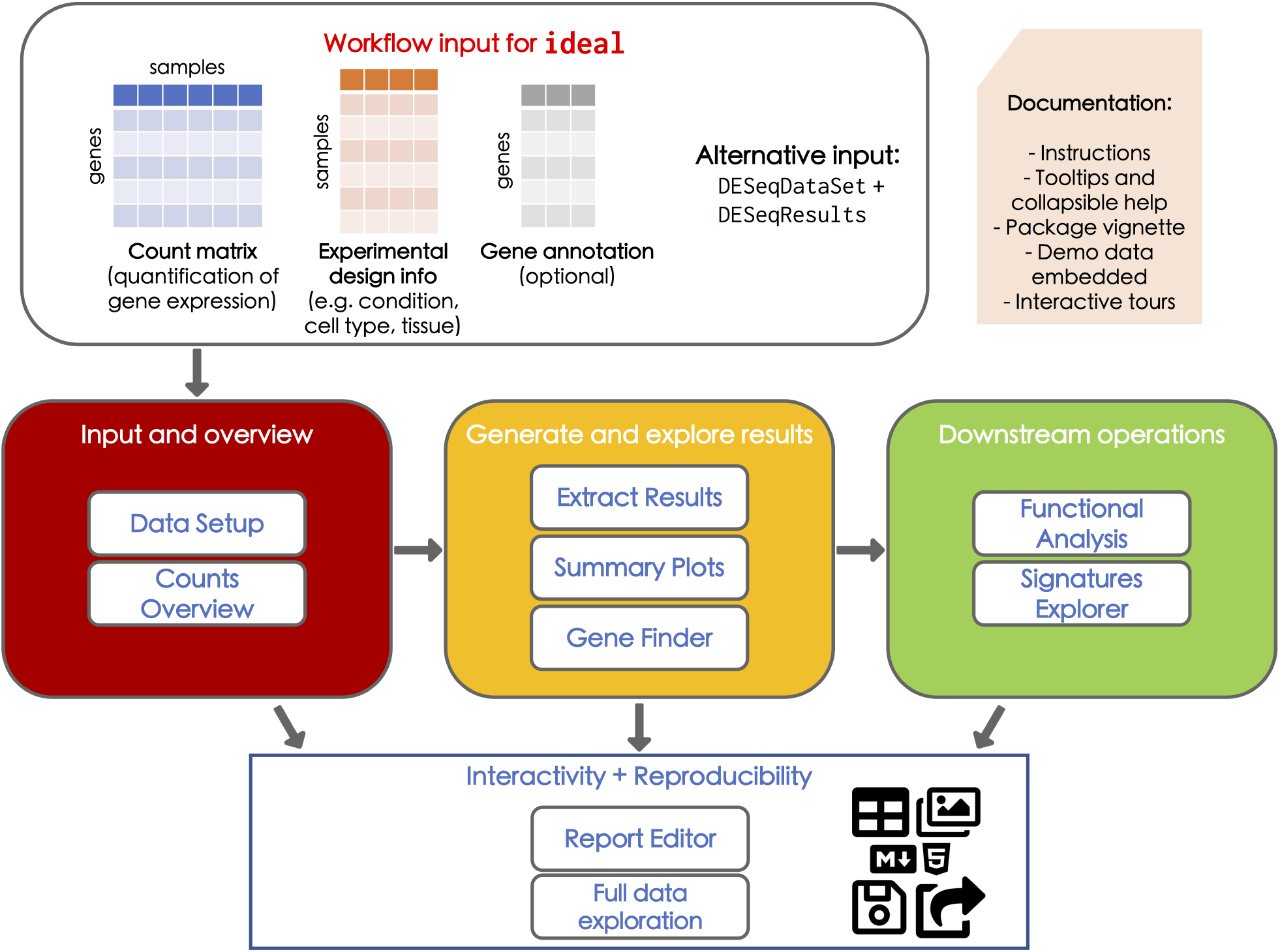
Overview of the ideal workflow. Top section: The typical analysis with `ideal` starts by providing the count matrix for the samples of interest, together with the corresponding experimental design information. The optional gene annotation information can also be retrieved at runtime. The combination of a `DESeqDataSet` and a `DESeqResults` objects can be given as an alternate input. Package documentation includes tooltips, collapsible help elements, and instructions in the app. Together with the vignette as a detailed reference, the interactive tours guide users through the fundamental components in each step, coupled to the embedded demo dataset. Middle section: The interactive session spans from the overview on the provided input, to the generation of differential expression analysis results and their visualization, while supporting downstream operations such as functional analysis, to assist in the interpretation of the data. Bottom section: All the generated output elements can be downloaded (images, tables), as well as exported in form of a R Markdown/HTML report, a document that guarantees reproducible analyses and can be readily shared or stored. (Icons contained in this figure are contained in the collections released by Font Awesome under the CC BY 4.0 license)

The `base` and `ggplot2` (Wickham, 2016) graphics systems are used to generate static visualizations, enabling interactions by brushing or clicking on them in the Shiny framework. Interactive heatmaps are generated with the `d3heatmap` (Cheng and Galili, 2018) package, and tables are displayed as interactive objects for efficient navigation via the `DT` package (Xie, 2018).

We provide an R Markdown template for a complete DE analysis together with the package, and users can customize its contents by editing or adding chunks in the embedded editor (based on the `shinyAce` package (Nijs *et al.*, 2018)). Combining this object with the current status of the reactive widgets in the main tabs of the application, an HTML report is generated for preview at runtime, and can later be exported, shared with colleagues, or simply stored (Figure 1, bottom section).

`ideal` has been tested on macOS, Linux, and Windows. It can be downloaded from the Bioconductor project page (http://bioconductor.org/packages/ideal/), and its development version can be found at https://github.com/federicomarini/ideal/. Alternatively, `ideal` is also provided as a Bioconda recipe (Grüning *et al.*, 2018), simplifying the installation procedure in isolated software environments e.g. in combined use with Snakemake (Köster and Rahmann, 2012), with binaries available at https://anaconda.org/bioconda/bioconductor-ideal.

Since `ideal` is normally installed on local systems, its speed and performance will vary depending on the hardware specifications available. In our experience, a typical modern laptop or workstation with at least 8 GB RAM is sufficient to run `ideal` on a variety of datasets. The desired depth of exploration after performing the backbone of the DE analysis is the main factor influencing the time required for completing a session with `ideal`, after familiarizing with its interface, e.g. by following the introductory tours on the demo dataset included.

The functionality of the `ideal` package is extensively described in the package vignette, regularly generated via the Bioconductor build system, and also embedded in the Welcome tab. Documentation for each function is provided, with examples rendered at the Github project page https://federicomarini.github.io/ideal/, generated with the `pkgdown` package (Wickham and Hesselberth, 2018).

### Typical usage workflow

During the typical usage session of `ideal`, users need to provide (or upload) two essential components: (1) a gene-level count matrix (`countmatrix`), a common intermediate result after quantifying the expression measures in widely used workflows ((Anders *et al.*, 2013; Pertea *et al.*, 2016; Love *et al.*, 2018)), and (2), the metadata table (`expdesign`) with the experimental variables for the samples of interest, as illustrated in the top panel of Figure 1. `ideal` can accept any tabular text files, and uses simple heuristics to detect the delimiter used to separate the distinct fields; a preview on the uploaded files is shown in the collapsible boxes in the Step 1 of the *Data Setup* panel (Figure 2a). A modal dialog informs the users about the formatting expected in the input files, with matched sample names and gene identifiers specified as column or row names. We strongly advise to perform a thorough exploratory data analysis on the input high-dimensional data, as this is a fundamental requirement in each rigorous analysis workflow. Users can refer to the `pcaExplorer` package for this purpose if an interactive approach is desired.

**Figure 2:**
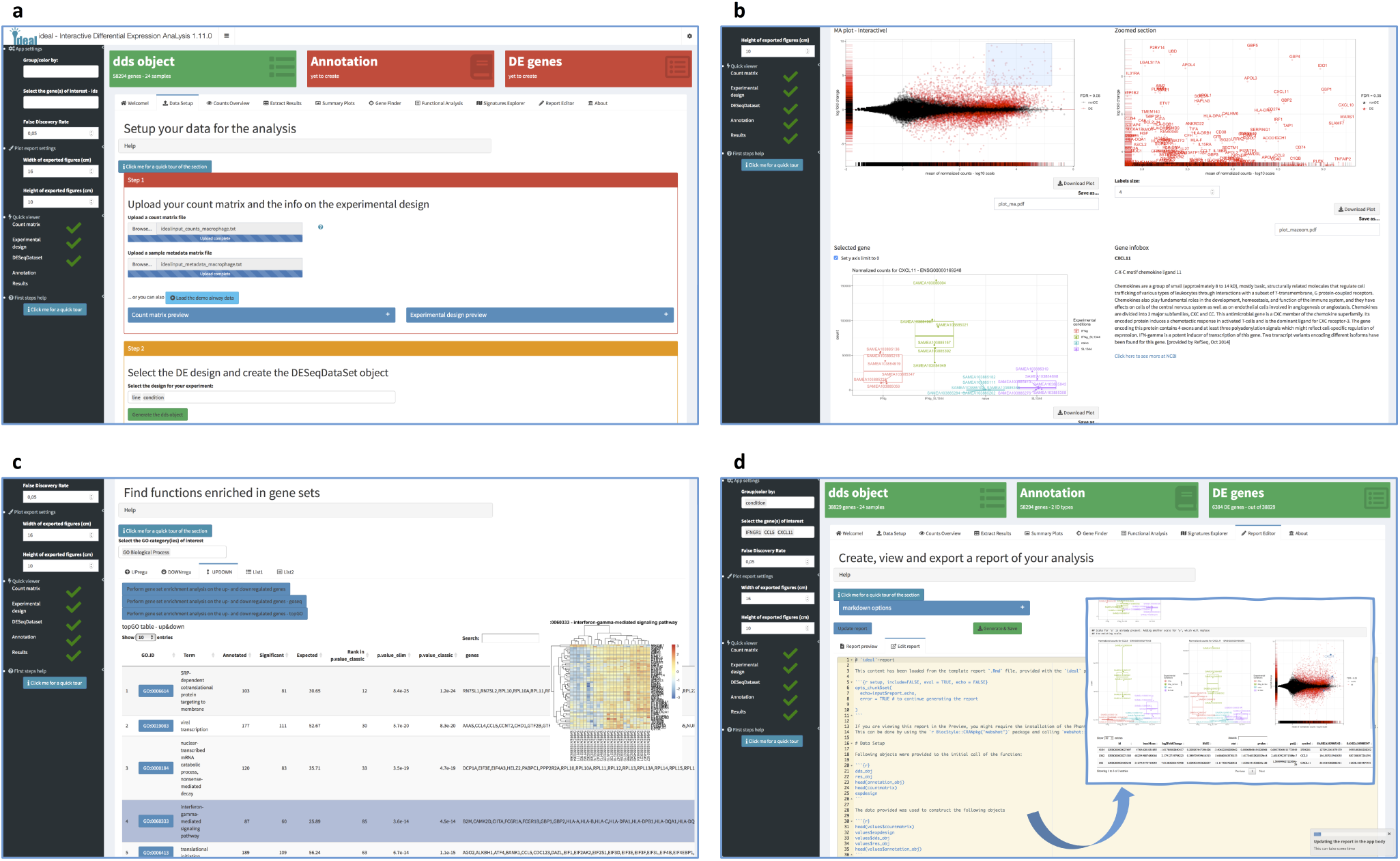
Selected screenshots of the ideal application. **a** Data Setup panel, after uploading the expression matrix and experimental design information, here displayed with the macrophage dataset illustrated as use case in Additional file 2. The setup steps appear consecutively once the required input elements and information are provided by the user. **b** Overview of the Summary Plots panel, with an MA plot displaying expression change (as log_2_ Fold Change) versus average expression level. Upon interaction via brushing, a zoomed plot appears, where features are labeled to facilitate further exploration, e.g. by displaying expression plots and information fetched from the NCBI Entrez database. **c** Results from the Functional Analysis panel, with an interactive table for the enriched Biological Processes. When clicking on a row of interest, a signature heatmap is shown to better show the behavior of the features annotated to the particular Gene Ontology term. **d** The Report Editor panel, accessed after computing all required objects (as shown by the green value boxes in the upper part of the subfigure), contains a text editor which displays the default comprehensive template provided with the package – which can additionally be edited by the user. In the framed content, a screenshot of the rendered report is included, e.g. with an annotated MA plot highlighting a set of specified genes.

In the context of differential expression testing, the design argument has also to be specified, and this is normally a subset of the variables in the experimental metadata, which constitute the main source of information when submitting the data to a public repository such as the NCBI Gene Expression Omnibus (National Center for Biotechnology Information, https://www.ncbi.nlm.nih.gov/geo/). All other parameters (the corresponding `DESeqDataSet`, `DESeqResults`, and a data frame containing matched identifiers for the features of interest) for the `ideal()` function can be also constructed manually on the command line and provided optionally, otherwise they will be computed at runtime.

For demonstration purposes, we include a primary human airway smooth muscle cell lines dataset (Himes *et al.*, 2014), which can be loaded in the *Data Setup* tab. For each module in the main application, `ideal` gives a text introduction to the typical operations, and then encourages the user to perform these in a guided manner by following the provided `rintrojs` tours, which can be started by clicking on a button. Descriptions of the user interface elements are anchored to the widgets themselves, and are highlighted in sequence while the interaction with them is enabled.

When the analysis session is terminated, binary RData objects and environments in the R session can store the exported reactive values. While all result files and figures generated and displayed in the user interface can also be saved locally with simple mouse clicks, the generation of a full interactive HTML report is the intended concluding step. This report is created by combining the values of reactive elements with the provided template, which can be extended by experienced users. Such a literate programming approach (conceived by (Knuth, 1984) and perfected in the `knitr` package) is one of the preferred methods to ensure the technical reproducibility of computational analyses (Sandve *et al.*, 2013; Stodden and Miguez, 2014).

Additionally, users can continue exploring interactively the exported objects, if some representations are not included in `ideal` directly. A flexible interface to do so is represented by the `iSEE` Bioconductor package (Rue-Albrecht *et al.*, 2018), which also fully tracks the code of the generated outputs, and we support this with a dedicated export function to a `SummarizedExperiment` object, with annotated `rowData` and `colData` slots filled with the results of the differential expression analysis.

### Deploying ideal on a Shiny server

While we anticipate that the `ideal` package will typically be installed on local machines, it can also be deployed as a web application on a Shiny server, simplifying the workflow for users who want to analyze and explore their data without installing software. Deployment of an instance shared among lab members of the same research group is an exemplary use case; our proposal also supports protected instances behind institutional firewalls, e.g. if sensitive patient data is to be handled.

We describe the full procedure to set up `ideal` on a server and document the required steps in the GitHub repository https://github.com/federicomarini/ideal_serveredition, which can be particularly useful for bioinformaticians or IT-system administrators. Following this approach, a publicly available instance has been created and is accessible at http://shiny.imbei.uni-mainz.de:3838/ideal for demonstration purposes, where users can either explore the `airway` dataset or upload their own data.

## Results

The functionality of `ideal` is described in the next sections, and is illustrated in detail for the analysis of a human RNA-seq data of macrophage immune stimulation (published in (Alasoo *et al.*, 2018)) in Additional file 2 (complete use case as HTML document structured like a vignette, with text, code, and output chunks).

### Data input and overview

The setup for the data analysis is carried out in the *Data Setup* panel (Figure 2a). To guide the user through the mandatory steps without an exceeding burden of interface elements, we designed this tab in a compact way, with boxes encapsulating related widgets, appearing consecutively once the upstream actions are completed. One of the fundamental data structures for the `ideal` app is a `DESeqDataSet` object, used in the workflow based on the `DESeq2` package (Anders *et al.*, 2013). This is complemented by an optional annotation object, i.e. a simple data frame where different key types (e.g. ENTREZ, ENSEMBL, HGNC-based gene symbols) are matched to the identifiers for the features of interest. While this is not mandatory, it is recommended as some of the package functionality relies on the interconversion across such identifiers; `ideal` suggests the corresponding orgDb Bioconductor packages and makes it immediate to create such an object directly at runtime. Once the initial selections are finalized, the `DESeq()` command runs the necessary steps of the pipeline, displaying a textual summary and a mean-dispersion plot as diagnostic tools.

A first overview on the dataset, including a set of basic summary statistics on the expressed genes, as well as the (log transformed) normalized values can be retrieved in the *Counts Overview* tab, together with pairwise scatter plots of the values. Thresholds can be introduced to subset the original dataset by keeping only genes with robust expression levels, either based on the detection in at least one sample, or on the average normalized value.

### Generating and exploring the results for Differential Expression analysis

The *Extract Results* tab provides the functionality to generate the other fundamental data structure, namely the `DESeqResults` object. After setting the FDR threshold in the sidebar, users are prompted to define the contrast of interest for their data, selecting one of the experimental factors included in the design. When the factor of interest has three or more levels available (e.g. the cell type in the `airway` demonstration dataset), the likelihood ratio test can be used instead of the Wald test, to allow for an ANOVA-like analysis across groups.

Further refinements to the results can be obtained by activating independent filtering (Bourgon *et al.*, 2010), or selecting the more powerful Independent Hypothesis Weighting (IHW) framework (Ignatiadis *et al.*, 2016), to ameliorate the multiple testing issue by incorporating an informative covariate, e.g. the mean gene expression (Korthauer *et al.*, 2019). Shrinkage of the effect sizes is also optionally performed on the log fold change estimates, to reflect the higher levels of uncertainty for lowly expressed genes. Interactive tables for the results are shown, with embedded links to the ENSEMBL browser and to the NCBI Gene portal to facilitate deeper exploration of shortlisted genes. Moreover, a number of diagnostic plots are generated, including histograms for unadjusted p-values, also using small multiples to stratify them on different mean expression value classes, a Schweder-Spjøtvoll plot (Schweder and Spjøtvoll, 1982), and a histogram of the estimated log fold changes.

More visualizations are included in the *Summary Plots* tab (Figure 2b), where users can zoom in the MA plot (M, log_2_ fold change vs A, average expression value) representation by brushing an area on the element. From the magnified subset, by clicking close to a selected gene, it is possible to obtain a gene expression boxplot (with the individual jittered observations superimposed), together with an info-box with details retrieved from the NCBI resource portal (Agarwala *et al.*, 2017). Heatmaps (both static and interactive) and volcano plots (log fold change vs log10 of the p-value) deliver alternative views of the underlying result table, or interesting subsets of it.

Iterations oriented towards exploring a set of features of interest are made easier by the *Gene Finder* tab. Genes can be shortlisted on the fly, adding them from the sidebar selectize widget. For each feature of interest, a plot comparing the normalized values is displayed, and these are included in an annotated MA plot, where the selected subsets are highlighted on the plot and their values are shown in a corresponding table. Alternatively, a gene list can be uploaded directly as text file to obtain the same output, with the ease of providing entire sets of genes (e.g. a file with all cytokines, or a curated list of genes affected by a particular transcription factor) in one step.

### Putting results into biological context

Many times it is challenging to make sense out of a carefully derived table of DE results, since it is not straightforward to identify the common biological themes that might be underlying the observed phenotypes. `ideal` offers different means to help researchers in meaningfully interpreting their RNA-seq data. The *Functional Analysis* panel offers three alternatives for gene set overrepresentation analysis, relying on `topGO` (Alexa *et al.*, 2006), `goseq` (Young *et al.*, 2010), or the `goana()` function in the `limma` package (Ritchie *et al.*, 2015); users can perform the enrichment tests on genes that are significantly differentially regulated, either split by direction of expression change, or combined in one list (Figure 2c). Additionally, users can upload up to two custom lists of genes, which can be compared to the one derived from the result object, in order to detect significant overlaps among the sets of interest, which can be represented via Venn diagrams or UpSet plots (Lex *et al.*, 2014).

The Gene Ontology (GO, (Ashburner *et al.*, 2000)) terms enriched in each list can be interactively displayed, with links to the AmiGO database (http://amigo.geneontology.org/), as well as heatmaps displaying the expression values for all the DE genes annotated to a particular signature.

Expanding on this functionality, `ideal` provides a *Signatures Explorer* panel, where a signature heatmap can be generated for any gene set provided in the gmt format, common to many sources of curated databases (MSigDB, WikiPathways). Conversion between identifier types is guided in the user interface, and so is the aspect of the final heatmap, where rows and columns can be clustered to better display existing patterns in the data, or transformations (such as mean centering or row standardization) help to bring the feature expression levels to a similar scale, for a better display in the final output.

### Generating reproducible and transparent results

The focus in the development of `ideal` was on combining interactivity and reproducibility of the analysis. Therefore, we implemented the *Report Editor* as the toolset for enabling reproducible reports in the DE analysis step (Figure 2d). The predefined template, embedded in the package, fetches the values of the reactive elements and the input widgets, thus capturing a snapshot of the ongoing session. Text, code, and results are all combined in an interactive HTML report, which can be previewed in the app, or subsequently exported.

This functionality is particularly appealing for less experienced users due to its automated simplicity, but experienced users can also take full benefit of it, by expanding the R Markdown document by adding or editing specific chunks of code.

State saving, activated by the buttons in the task menu in the header, stores the content of the current session into binary data objects or environments accessible from the global workspace.

As an additional feature, we leverage the flexibility of `iSEE`, the interactive Summarized-Experiment Explorer, another tool which fully supports intuitive and reproducible analyses, by assembling a serialized rds object that can be directly fed to the main `iSEE()` function for bespoke visualizations.

## Discussion

The guiding principle for the development of our package `ideal` was the effective combination of usability and reproducibility, applied to one of the most widely adopted workflows in transcriptomics, i.e. the analysis of differentially expressed genes, followed by the down-stream analyses based on functional enrichment among the subset of detected features.

Several software packages have been developed to operate on this tabular-like summarized expression data, or on formats which might derive from their results, and a comprehensive comparison of their features is presented in Additional file 1: Table S1. Notably, these tools differ by their set of included features (ranging from first exploration to downstream analysis steps), implementation (with R/Shiny, python, and JavaScript as main choices), format of distribution (packages, local web app, webserver), and ease of implementation in existing pipelines (e.g. by leveraging widely adopted class structures, or requiring and providing text files for portability across systems). The comparison with other tools is also available online (https://federicomarini.github.io/ideal_supplement), linked to a Google Sheet where the individual characteristics of each tool will be updated, in order to provide a tool for users who might be seeking advice on which solution to adapt for their needs (accessible at https://docs.google.com/spreadsheets/d/167XV0w18P0FSld1dt6owN4C2Esxl5FU2QTo4D-wclz0/edit?usp=sharing).

Our proposal complements the existing `pcaExplorer` package, where the main exploratory data analysis steps are performed, and provides a platform for performing a complete differential expression analysis with the ease of interactivity, accompanied by a number of diagnostic plots, often overseen in other software tools. The combination of interactivity with reproducibility (Figure 1, bottom panel) is an essential aspect to consider for generating robust and transparent analyses, substantiated by code which can also be used for didactic purposes, learning the best current practices with the state-of-the-art methods included in our package.

`ideal` fully supports widely used standard classes from Bioconductor, and thus allows seamless integration with many R packages for further downstream processing and within existing data analysis pipelines, while also benefiting from a thriving community of developers. `ideal` itself is part of Bioconductor, and thus is integrated into a build system that continuously checks all of its components and their interoperability, guaranteeing that the available set of features is correctly interfacing with the latest version of the package dependencies. Notably, the Bioconductor project enforces a number of best practices to enhance the usability of its components, with both internal and external documentation (for the individual functions and as complete tutorials, in form of vignettes), as well as providing unit test sets to ensure the software is working as expected: `ideal` adheres to these guidelines, which can be essential to define robust software (Taschuk and Wilson, 2017) that can be adopted by a wide range of users.

A possible use case for deploying `ideal` is tightly related to data and result sharing. Distributing data in raw and processed format, together with a set of results, is becoming a practice that enables efficient data mining and can help ensure their reproducibility and reusability (Wilkinson *et al.*, 2016). Stemming from a close collaboration with life and medical scientists, our tool allows researchers to share their work with other interested parties, starting from the operations during the collaboration phase, and continuing after publication, where broader audiences can effectively digest contents as they are presented from the authors.

The faster turnover in generating insights, thanks to the accessible interface and the multiple outputs, constitutes a significant advantage for reducing the time to new results, or alternatively for re-analyzing publicly available RNA-seq datasets. `ideal` provides a platform to facilitate discoveries in a standardized way, which at the same time improves the transparency and the reproducibility of the analyses. Indeed, one possible use case that we envision is the submission of a comprehensive notebook/report as Supplementary Material for a manuscript, so that the results are presented in a transparent manner, thus facilitating the contribution of reviewers, as well as the re-usability of analysis code. A rich technical description of parameters and software used would also greatly facilitate the writing of the “Material and Methods” sections, in a way that fully captures steps, parameters, code, and software versions.

## Conclusion

The infrastructure provided by the `ideal` R/Bioconductor package delivers a web browser application that guarantees ease of use through interactivity and a dynamic user interface, together with reproducible research, for the essential step of differential expression investigation in RNA-seq analysis. The combination of these two features is a key factor for efficient, quick, and robust extraction of information, while leveraging the facilities available in the Bioconductor project in terms of classes and statistical methods.

The wealth of information that can be extracted while running the app may play a critical role when choosing the tools to adopt in a project. Still, to ensure the proper interpretation of the output results, the interaction of wet-lab scientists with collaborators with additional bioinformatics/biostatistics expertise is essential. The design choices for `ideal` aim at making this communication as robust and easy as possible, possibly defining this tool as the *ideal* way of approaching this step.

Following the criteria used in our previous overview on RNA-seq analysis interfaces (Poplawski *et al.*, 2016), our package reaches out to the life/medical scientist, being simple to install and use, based on robust statistical methods, and offering multiple levels of documentation. `ideal` allows scientists to easily take control of the analysis of RNA-seq data, while providing an accessible framework for reproducible research, which can be extended according to the user’s needs.

## Supporting information

Additional File 1 – Table S1, Comparison of software

Additional File 2 – Complete use case for the ideal package (zipped HTML document)

## Additional information

### Availability and requirements

**Project name:** ideal

**Project home page:** http://bioconductor.org/packages/ideal/ (release) and https://github.com/federicomarini/ideal/ (development version)

**Archived version:** https://doi.org/10.5281/zenodo.3601634, package source as gzipped tar archive of the version reported in this article

**Project documentation:** rendered at https://federicomarini.github.io/ideal/

**Operating systems:** Linux, Mac OS, Windows

**Programming language:** R

**Other requirements:** R 3.4 or higher, Bioconductor 3.5 or higher

**License:** MIT

**Any restrictions to use by non-academics:** none.

## Abbreviations

ANOVA: Analysis of Variance
DE: Differential Expression
FDR: False Discovery Rate
GO: Gene Ontology
HGNC: HUGO (Human Genome Organisation) Gene Nomenclature Committee
IHW: Independent Hypothesis Weighting
MA plot: M (log ratio) vs A (mean average) plot
MSigDB: Molecular Signatures Database
NCBI: National Center for Biotechnology Information
RNA-seq: RNA sequencing

## Acknowledgements

We thank Sebastian Schubert and Carina Santos of the Ruf lab (CTH Mainz) for fruitful discussions and their feedback as early adopters of the `ideal` package, as well as the users’ community for their helpful suggestions. We also thank Charlotte Soneson (FMI, Basel), Miguel Andrade, Wolfram Ruf, Franziska Härtner, Gerrit Toenges, and Konstantin Strauch for their helpful comments on the manuscript.

## Availability of data and materials

Data used in the described use cases (as demo dataset and in Additional file 2) is available from the following articles:

- The airway smooth muscle cell RNA-seq is included in PubMed ID: 24926665 (https://doi.org/10.1371/journal.pone.0099625). GEO entry: GSE52778, accessed from the Bioconductor experiment package `airway` (http://bioconductor.org/packages/airway/, version 1.7.0).
- The data set on the macrophage immune stimulation is included in PubMed ID: 29379200 (https://doi.org/10.1038/s41588-018-0046-7). Dataset deposited at the ENA (ERP020977, project id: PRJEB18997) and accessed from the Bioconductor experiment package `macrophage` package (http://bioconductor.org/packages/macrophage/, version 1.3.1)

The `ideal` package can be downloaded from its Bioconductor page http://bioconductor.org/packages/ideal/ or the GitHub development page https://github.com/federicomarini/ideal/. `ideal` is also provided as a recipe in Bioconda (https://anaconda.org/bioconda/bioconductor-ideal).

## Additional Files

- **Additional file 1 — Table S1 (AddFile1_ideal_supplement.xlsx, .xlsx file)** Comparison of software for analyzing interactively RNA-Seq data, including link to the related publications (if available) and to the source code repositories. Evaluation criteria are included in the dedicated sheet. The information contained in this table are also available online at https://federicomarini.github.io/ideal_supplement, displaying the content of the `ideal_supplement` Google Sheet, accessible at https://docs.google.com/spreadsheets/d/167XV0w18P0FSld1dt6owN4C2Esxl5FU2QTo4D-wclz0/edit?usp=sharing).
- **Additional file 2 (AddFile2_usecase_ideal.html.zip, compressed HTML document)** Complete use case for the `ideal` package, based on the `macrophage` immune stimulation dataset (Interferon Gamma treatment vs naive cells).

## Authors’ contributions

F.M. – Conceptualization, Methodology, Software Implementation, Writing – Original Draft Preparation, Writing – Review & Editing

J.L. – Software Implementation, Writing – Review & Editing

H.B. – Supervision, Writing – Review & Editing

All authors read and approved the final version of the manuscript.

## Ethics approval and consent to participate

Not applicable.

## Consent for publication

Not applicable.

## Bibliography

Agarwala, R., Barrett, T., Beck, J., Benson, D. A., Bollin, C., Bolton, E., Bourexis, D., Brister, J. R., Bryant, S. H., Canese, K., Charowhas, C., Clark, K., DiCuccio, M., Dondoshansky, I., Feolo, M., Funk, K., Geer, L. Y., Gorelenkov, V., Hlavina, W., Hoeppner, M., Holmes, B., Johnson, M., Khotomlianski, V., Kimchi, A., Kimelman, M., Kitts, P., Klimke, W., Krasnov, S., Kuznetsov, A., Landrum, M. J., Landsman, D., Lee, J. M., Lipman, D. J., Lu, Z., Madden, T. L., Madej, T., Marchler-Bauer, A., Karsch-Mizrachi, I., Murphy, T., Orris, R., Ostell, J., O’Sullivan, C., Palanigobu, V., Panchenko, A. R., Phan, L., Pruitt, K. D., Rodarmer, K., Rubinstein, W., Sayers, E. W., Schneider, V., Schoch, C. L., Schuler, G. D., Sherry, S. T., Sirotkin, K., Siyan, K., Slotta, D., Soboleva, A., Soussov, V., Starchenko, G., Tatusova, T. A., Todorov, K., Trawick, B. W., Vakatov, D., Wang, Y., Ward, M., Wilbur, W. J., Yaschenko, E., and Zbicz, K. (2017). Database Resources of the National Center for Biotechnology Information. Nucleic Acids Research, 45(D1), D12–D17.

Alasoo, K., Rodrigues, J., Mukhopadhyay, S., Knights, A. J., Mann, A. L., Kundu, K., Hale, C., Dougan, G., and Gaffney, D. J. (2018). Shared genetic effects on chromatin and gene expression indicate a role for enhancer priming in immune response. Nature Genetics, 50(3), 424–431.

Alexa, A., Rahnenführer, J., and Lengauer, T. (2006). Improved scoring of functional groups from gene expression data by decorrelating GO graph structure. Bioinformatics, 22(13), 1600–1607.

Allaire, J., Xie, Y., McPherson, J., Luraschi, J., Ushey, K., Atkins, A., Wickham, H., Cheng, J., and Chang, W. (2018). rmarkdown: Dynamic Documents for R. R package version 1.10.

Amezquita, R. A., Carey, V. J., Carpp, L. N., Geistlinger, L., Lun, A. T., Marini, F., Rue-Albrecht, K., Risso, D., Soneson, C., Waldron, L., Pages, H., Smith, M., Huber, W., Morgan, M., Gottardo, R., and Hicks, S. C. (2019). Orchestrating Single-Cell Analysis with Bioconductor. bioRxiv, page 590562.

Anders, S., McCarthy, D. J., Chen, Y., Okoniewski, M., Smyth, G. K., Huber, W., and Robinson, M. D. (2013). Count-based differential expression analysis of RNA sequencing data using R and Bioconductor. Nature Protocols, 8(9), 1765–1786.

Anders, S., Pyl, P. T., and Huber, W. (2015). HTSeq–a Python framework to work with high-throughput sequencing data. Bioinformatics, 31(2), 166–169.

Ashburner, M., Ball, C. A., Blake, J. A., Botstein, D., Butler, H., Cherry, J. M., Davis, A. P., Dolinski, K., Dwight, S. S., Eppig, J. T., Harris, M. A., Hill, D. P., Issel-Tarver, L., Kasarskis, A., Lewis, S., Matese, J. C., Richardson, J. E., Ringwald, M., Rubin, G. M., and Sherlock, G. (2000). Gene Ontology: tool for the unification of biology. Nature Genetics, 25(1), 25–29.

Bailey, E. (2015). shinyBS: Twitter Bootstrap Components for Shiny. R package version 0.61.

Beigh, M. (2016). Next-Generation Sequencing: The Translational Medicine Approach from “Bench to Bedside to Population”. Medicines, 3(2), 14.

Bourgon, R., Gentleman, R., and Huber, W. (2010). Independent filtering increases detection power for high-throughput experiments. Proceedings of the National Academy of Sciences, 107(21), 9546–9551.

Bray, N. L., Pimentel, H., Melsted, P., and Pachter, L. (2016). Near-optimal probabilistic RNA-seq quantification. Nature Biotechnology, 34(5), 525–527.

Chang, W. and Borges Ribeiro, B. (2018). shinydashboard: Create Dashboards with ‘Shiny’. R package version 0.7.0.

Chang, W., Cheng, J., Allaire, J. J., Xie, Y., and McPherson, J. (2016). shiny: Web Application Framework for R.

Cheng, J. and Galili, T. (2018). d3heatmap: Interactive Heat Maps Using ‘htmlwidgets’ and ‘D3.js’. R package version 0.6.1.2.

Choi, K. and Ratner, N. (2019). iGEAK: an interactive gene expression analysis kit for seamless workflow using the R/shiny platform. BMC Genomics, 20(1), 177.

Conesa, A., Madrigal, P., Tarazona, S., Gomez-Cabrero, D., Cervera, A., McPherson, A., Szcześniak, M. W., Gaffney, D. J., Elo, L. L., Zhang, X., and Mortazavi, A. (2016). A survey of best practices for RNA-seq data analysis. Genome Biology, 17(1), 13.

Dobin, A., Davis, C. A., Schlesinger, F., Drenkow, J., Zaleski, C., Jha, S., Batut, P., Chaisson, M., and Gingeras, T. R. (2013). STAR: ultrafast universal RNA-seq aligner. Bioinformatics, 29(1), 15–21.

Eglen, S. J., Marwick, B., Halchenko, Y. O., Hanke, M., Sufi, S., Gleeson, P., Silver, R. A., Davison, A. P., Lanyon, L., Abrams, M., Wachtler, T., Willshaw, D. J., Pouzat, C., and Poline, J.-B. (2017). Toward standard practices for sharing computer code and programs in neuroscience. Nature Neuroscience, 20(6), 770–773.

Froussios, K., Schurch, N. J., Mackinnon, K., Gierliński, M., Duc, C., Simpson, G. G., and Barton, G. J. (2019). How well do RNA-Seq differential gene expression tools perform in a complex eukaryote? A case study in Arabidopsis thaliana. Bioinformatics, page 090753.

Ganz, C. (2016). rintrojs: A Wrapper for the Intro.js Library. The Journal of Open Source Software, 1(6), 2016.

Gardeux, V., David, F. P., Shajkofci, A., Schwalie, P. C., and Deplancke, B. (2017). ASAP: A web-based platform for the analysis and interactive visualization of single-cell RNA-seq data. Bioinformatics, 33(19), 3123–3125.

Ge, S. X., Son, E. W., and Yao, R. (2018). iDEP: an integrated web application for differential expression and pathway analysis of RNA-Seq data. BMC Bioinformatics, 19(1), 534.

Gentleman, R. C., Carey, V. J., Bates, D. M., Bolstad, B., Dettling, M., Dudoit, S., Ellis, B., Gautier, L., Ge, Y., Gentry, J., Hornik, K., Hothorn, T., Huber, W., Iacus, S., Irizarry, R., Leisch, F., Li, C., Maechler, M., Rossini, A. J., Sawitzki, G., Smith, C., Smyth, G., Tierney, L., Yang, J. Y. H., and Zhang, J. (2004). Bioconductor: open software development for computational biology and bioinformatics. Genome Biology, 5(10), R80.

Grüning, B., Dale, R., Sjödin, A., Chapman, B. A., Rowe, J., Tomkins-Tinch, C. H., Valieris, R., and Köster, J. (2018). Bioconda: Sustainable and comprehensive software distribution for the life sciences. Nature Methods, 15(7), 475–476.

Harshbarger, J., Kratz, A., and Carninci, P. (2017). DEIVA: a web application for interactive visual analysis of differential gene expression profiles. BMC Genomics, 18(1), 47.

Himes, B. E., Jiang, X., Wagner, P., Hu, R., Wang, Q., Klanderman, B., Whitaker, R. M., Duan, Q., Lasky-Su, J., Nikolos, C., Jester, W., Johnson, M., Panettieri, R. a., Tantisira, K. G., Weiss, S. T., and Lu, Q. (2014). RNA-Seq Transcriptome Profiling Identifies CRISPLD2 as a Glucocorticoid Responsive Gene that Modulates Cytokine Function in Airway Smooth Muscle Cells. PLoS ONE, 9(6), e99625.

Huber, W., Carey, V. J., Gentleman, R., Anders, S., Carlson, M., Carvalho, B. S., Bravo, H. C., Davis, S., Gatto, L., Girke, T., Gottardo, R., Hahne, F., Hansen, K. D., Irizarry, R. a., Lawrence, M., Love, M. I., MacDonald, J., Obenchain, V., Oleś, A. K., Pagès, H., Reyes, A., Shannon, P., Smyth, G. K., Tenenbaum, D., Waldron, L., and Morgan, M. (2015). Orchestrating high-throughput genomic analysis with Bioconductor. Nature Methods, 12(2), 115–121.

Ignatiadis, N., Klaus, B., Zaugg, J. B., and Huber, W. (2016). Data-driven hypothesis weighting increases detection power in genome-scale multiple testing. Nature Methods, 13(7), 577–580.

Knuth, D. E. (1984). Literate Programming. The Computer Journal, 27(2), 97–111.

Korthauer, K., Kimes, P. K., Duvallet, C., Reyes, A., Subramanian, A., Teng, M., Shukla, C., Alm, E. J., and Hicks, S. C. (2019). A practical guide to methods controlling false discoveries in computational biology. Genome Biology, 20(1), 118.

Köster, J. and Rahmann, S. (2012). Snakemake–a scalable bioinformatics workflow engine. Bioinformatics, 28(19), 2520–2522.

Kucukural, A., Yukselen, O., Ozata, D. M., Moore, M. J., and Garber, M. (2019). DEBrowser: interactive differential expression analysis and visualization tool for count data. BMC Genomics, 20(1), 6.

Lex, A., Gehlenborg, N., Strobelt, H., Vuillemot, R., and Pfister, H. (2014). UpSet: Visualization of intersecting sets. IEEE Transactions on Visualization and Computer Graphics, 20(12), 1983–1992.

Li, Y. and Andrade, J. (2017). DEApp: An interactive web interface for differential expression analysis of next generation sequence data. Source Code for Biology and Medicine, 12(1), 10–13.

Liao, Y., Smyth, G. K., and Shi, W. (2014). featureCounts: an efficient general purpose program for assigning sequence reads to genomic features. Bioinformatics, 30(7), 923–930.

Lim, J. H., Lee, S. Y., and Kim, J. H. (2017). TRAPR: R Package for Statistical Analysis and Visualization of RNA-Seq Data. Genomics & Informatics, 15(1), 51.

Love, M. I., Huber, W., and Anders, S. (2014). Moderated estimation of fold change and dispersion for RNA-seq data with DESeq2. Genome Biology, 15(12), 550.

Love, M. I., Anders, S., Kim, V., and Huber, W. (2015). RNA-Seq workflow: gene-level exploratory analysis and differential expression. F1000Research, 4, 1070.

Love, M. I., Soneson, C., and Patro, R. (2018). Swimming downstream: statistical analysis of differential transcript usage following Salmon quantification, volume 7.

Marini, F. and Binder, H. (2016). Development of Applications for Interactive and Reproducible Research : a Case Study. Genomics and Computational Biology, 3(1), 1–4.

Marini, F. and Binder, H. (2019). pcaExplorer: an R/Bioconductor package for interacting with RNA-seq principal components. BMC Bioinformatics, 20(1), 331.

McCarthy, D. J., Chen, Y., and Smyth, G. K. (2012). Differential expression analysis of multifactor RNA-Seq experiments with respect to biological variation. Nucleic Acids Research, 40(10), 4288–4297.

McDermaid, A., Monier, B., Zhao, J., Liu, B., and Ma, Q. (2018). Interpretation of differential gene expression results of RNA-seq data: review and integration. Briefings in Bioinformatics, 00(April), 1–11.

McNutt, M. (2014). Journals unite for reproducibility. Science, 346(6210), 679–679.

Monier, B., McDermaid, A., Zhao, J., Fennell, A., and Ma, Q. (2018). IRIS-EDA: An integrated RNA-Seq interpretation system for gene expression data analysis. bioRxiv, page 283341.

Nelson, J. W., Sklenar, J., Barnes, A. P., and Minnier, J. (2016). The START App: a web-based RNAseq analysis and visualization resource. Bioinformatics, 33(3), btw624.

Nijs, V., Fang, F., Trestle Technology, LLC, and Allen, J. (2018). shinyAce: Ace Editor Bindings for Shiny. R package version 0.3.2.

Nosek, B. A., Alter, G., Banks, G. C., Borsboom, D., Bowman, S. D., Breckler, S. J., Buck, S., Chambers, C. D., Chin, G., Christensen, G., Contestabile, M., Dafoe, A., Eich, E., Freese, J., Glennerster, R., Goroff, D., Green, D. P., Hesse, B., Humphreys, M., Ishiyama, J., Karlan, D., Kraut, A., Lupia, A., Mabry, P., Madon, T., Malhotra, N., Mayo-Wilson, E., McNutt, M., Miguel, E., Paluck, E. L., Simonsohn, U., Soderberg, C., Spellman, B. A., Turitto, J., VandenBos, G., Vazire, S., Wagenmakers, E. J., Wilson, R., and Yarkoni, T. (2015). Promoting an open research culture. Science, 348(6242), 1422–1425.

Oshlack, A., Robinson, M. D., and Young, M. D. (2010). From RNA-seq reads to differential expression results. Genome Biology, 11(12), 220.

Patro, R., Duggal, G., Love, M. I., Irizarry, R. A., and Kingsford, C. (2017). Salmon provides fast and bias-aware quantification of transcript expression. Nature Methods.

Peng, R. D. (2011). Reproducible Research in Computational Science. Science, 334(6060), 1226–1227.

Perkel, J. M. (2018). Data visualization tools drive interactivity and reproducibility in online publishing. Nature, 554(7690), 133–134.

Pertea, M., Kim, D., Pertea, G. M., Leek, J. T., and Salzberg, S. L. (2016). Transcript-level expression analysis of RNA-seq experiments with HISAT, StringTie and Ballgown. Nature Protocols, 11(9), 1650–1667.

Poplawski, A., Marini, F., Hess, M., Zeller, T., Mazur, J., and Binder, H. (2016). Systematically evaluating interfaces for RNA-seq analysis from a life scientist perspective. Briefings in Bioinformatics, 17(2), 213–223.

Price, A., Caciula, A., Guo, C., Lee, B., Morrison, J., Rasmussen, A., Lipkin, W. I., and Jain, K. (2019). DEvis: an R package for aggregation and visualization of differential expression data. BMC Bioinformatics, pages 1–7.

Ritchie, M. E., Phipson, B., Wu, D., Hu, Y., Law, C. W., Shi, W., and Smyth, G. K. (2015). Limma powers differential expression analyses for RNA-sequencing and microarray studies. Nucleic Acids Research, 43(7), e47.

Rue-Albrecht, K., Marini, F., Soneson, C., and Lun, A. T. (2018). iSEE: Interactive SummarizedExperiment Explorer. F1000Research, 7(0), 741.

Sandve, G. K., Nekrutenko, A., Taylor, J., and Hovig, E. (2013). Ten Simple Rules for Reproducible Computational Research. PLoS Computational Biology, 9(10), e1003285.

Schultheis, H., Kuenne, C., Preussner, J., Wiegandt, R., Fust, A., Bentsen, M., and Looso, M. (2018). WIlsON: Web-based Interactive Omics VisualizatioN. Bioinformatics, 33(17), 2699–2705.

Schurch, N. J., Schofield, P., Gierliński, M., Cole, C., Sherstnev, A., Singh, V., Wrobel, N., Gharbi, K., Simpson, G. G., Owen-Hughes, T., Blaxter, M., and Barton, G. J. (2016). How many biological replicates are needed in an RNA-seq experiment and which differential expression tool should you use? RNA, 22(6), 839–851.

Schweder, T. and Spjøtvoll, E. (1982). Plots of P-values to evaluate many tests simultaneously. Biometrika, 69(3), 493–502.

Soneson, C., Love, M. I., and Robinson, M. D. (2015). Differential analyses for RNA-seq: transcript-level estimates improve gene-level inferences. F1000Research, 4(0), 1521.

Stodden, B. V., Mcnutt, M., Bailey, D. H., Deelman, E., Hanson, B., Heroux, M. A., Ioannidis, J. P. A., and Taufer, M. (2016). Enhancing reproducibility for computational methods. Science, 354(6317), 1240–1241.

Stodden, V. and Miguez, S. (2014). Best Practices for Computational Science: Software Infrastructure and Environments for Reproducible and Extensible Research. Journal of Open Research Software, 2(1), e21.

Su, S., Law, C. W., Ah-Cann, C., Asselin-Labat, M.-L., Blewitt, M. E., and Ritchie, M. E. (2017). Glimma: interactive graphics for gene expression analysis. Bioinformatics, 33(13), 2050–2052.

Su, W., Sun, J., Shimizu, K., and Kadota, K. (2019). TCC-GUI: a Shiny-based application for differential expression analysis of RNA-Seq count data. BMC Research Notes, 12(1), 133.

Taschuk, M. and Wilson, G. (2017). Ten simple rules for making research software more robust. PLOS Computational Biology, 13(4), e1005412.

Tintori, S. C., Golden, P., and Goldstein, B. (2020). Differential Expression Gene Explorer (DrEdGE): A tool for generating interactive online visualizations of gene expression datasets. Bioinformatics, 8(5), 55.

Van den Berge, K., Hembach, K. M., Soneson, C., Tiberi, S., Clement, L., Love, M. I., Patro, R., and Robinson, M. D. (2019). RNA Sequencing Data: Hitchhiker’s Guide to Expression Analysis. Annual Review of Biomedical Data Science, 2(1), 139–173.

Wang, Z., Gerstein, M., and Snyder, M. (2009). RNA-Seq: a revolutionary tool for transcriptomics. Nature Reviews Genetics, 10(1), 57–63.

Wickham, H. (2016). ggplot2: Elegant Graphics for Data Analysis. Springer-Verlag New York.

Wickham, H. and Hesselberth, J. (2018). pkgdown: Make Static HTML Documentation for a Package. R package version 1.1.0.

Wilkinson, M. D., Dumontier, M., Aalbersberg, I. J., Appleton, G., Axton, M., Baak, A., Blomberg, N., Boiten, J.-W., da Silva Santos, L. B., Bourne, P. E., Bouwman, J., Brookes, A. J., Clark, T., Crosas, M., Dillo, I., Dumon, O., Edmunds, S., Evelo, C. T., Finkers, R., Gonzalez-Beltran, A., Gray, A. J., Groth, P., Goble, C., Grethe, J. S., Heringa, J., ‘t Hoen, P. A., Hooft, R., Kuhn, T., Kok, R., Kok, J., Lusher, S. J., Martone, M. E., Mons, A., Packer, A. L., Persson, B., Rocca-Serra, P., Roos, M., van Schaik, R., Sansone, S.-A., Schultes, E., Sengstag, T., Slater, T., Strawn, G., Swertz, M. A., Thompson, M., van der Lei, J., van Mulligen, E., Velterop, J., Waagmeester, A., Wittenburg, P., Wolstencroft, K., Zhao, J., and Mons, B. (2016). The FAIR Guiding Principles for scientific data management and stewardship. Scientific Data, 3, 160018.

Xie, Y. (2018). DT: A Wrapper of the JavaScript Library ‘DataTables’. R package version 0.4.

Younesy, H., Möller, T., Lorincz, M. C., Karimi, M. M., and Jones, S. J. (2015). VisRseq: R-based visual framework for analysis of sequencing data. BMC Bioinformatics, 16(Suppl 11), S2.

Young, M. D., Wakefield, M. J., Smyth, G. K., and Oshlack, A. (2010). Gene ontology analysis for RNA-seq: accounting for selection bias. Genome Biology, 11(2), R14.

Zhu, Q., Fisher, S. A., Dueck, H., Middleton, S., Khaladkar, M., and Kim, J. (2018). PIVOT: platform for interactive analysis and visualization of transcriptomics data. BMC Bioinformatics, 19(1), 6.

