## Additional File 2 – Complete use case for the ideal package (zipped HTML document) for "ideal: an R/Bioconductor package for Interactive Differential Expression Analysis": AddFile2_usecase_ideal.html

A complete use case for the ideal package


Code 

- Show All Code
- Hide All Code
- Download Rmd

### A complete use case for the *ideal* package

###### Federico Marini

Center for Thrombosis and Hemostasis (CTH), Mainz;  
Institute of Medical Biostatistics, Epidemiology and Informatics (IMBEI), Mainz  


###### Jan Linke

Center for Thrombosis and Hemostasis (CTH), Mainz;  
Institute of Medical Biostatistics, Epidemiology and Informatics (IMBEI), Mainz  

###### Harald Binder

Institute of Medical Biometry and Statistics, Faculty of Medicine and Medical Center - University of Freiburg  

###### 6 January 2020

**Compiled date**: 2020-01-06

**Last edited**: 2020-01-06

**License**: MIT + file LICENSE

### 1 Introduction

This document contains a fully worked out use case for the *ideal* package, using the RNA-seq dataset from Alasoo, et al. “Shared genetic effects on chromatin and gene expression indicate a role for enhancer priming in immune response”, published in Nature Genetics, January 2018 (Alasoo et al. 2018)
doi:10.1038/s41588-018-0046-7.

The data is made available via the *macrophage* Bioconductor package, which contains the files output from the Salmon quantification (version 0.12.0, with Gencode v29 reference).

As for the experimental setting, the samples are available from 6 different donors, in 4 different conditions (naive, treated with Interferon gamma, with SL1344, or with a combination of Interferon gamma and SL1344).

**Note:** this document has been created coupled to the implementation of *ideal* at the moment of submission, in version 1.11.0, deposited at Zenodo []TODO LINK[].
Slight variations in the aspect and behavior will reflect the newer developments of the package.

Throughout this document, you will encounter small text boxes shaded in light blue: these will contain the essential information specific to the `macrophage` dataset.
You can follow these steps as a quick way of navigating the complete walkthrough.

### 2 Installation and setup

The following packages need to be installed to reproduce the analyses in this document:

```
install.packages("BiocManager")         # for installing Bioconductor
BiocManager::install(c("ideal",
                       "macrophage",
                       "DESeq2",
                       "org.Hs.eg.db",
                       "tximeta",
                       "clusterProfiler")
                     )
```

Loading the required packages:

```
library("macrophage")
library("magrittr")
library("SummarizedExperiment")
library("tximeta")
library("DESeq2")
library("AnnotationDbi")
library("org.Hs.eg.db")
library("pheatmap")
library("clusterProfiler")
library("knitr")
library("pcaExplorer")
library("ideal")
```

#### 2.1 Launching `ideal()`

The `ideal()` function has some essential parameters, and it is worth elaborating some details on each:

- `dds_obj` - a `DESeqDataSet` object. If not provided, then a `countmatrix` and a `expdesign` need to be provided. If none of the above is provided, it is possible to upload the data during the execution of the Shiny App.
- `res_obj` - a `DESeqResults` object. If not provided, it can be computed during the execution of the application. This is the basic class to store the result for the comparison of interest.
- `annotation_obj` - a `data.frame` object, with `row.names` as gene identifiers (e.g. ENSEMBL ids) and a column, `gene_name`, containing e.g. HGNC-based gene symbols. If not provided, it can be constructed during the execution via the `org.eg.XX.db` packages; this can be quite handy as these packages can work with a wide range of identifier types.
- `countmatrix` - a count matrix, with genes as rows and samples as columns. If not provided, it is possible to upload the data during the execution of the Shiny App.
- `expdesign` - a `data.frame` containing the info on the experimental covariates of each sample. If not provided, it is possible to upload the data during the execution of the Shiny App.
- `gene_signatures` - a list of vectors, one for each pathway/signature. This is for example the output of the `read_gmt` function, which takes as input files in the gmt format, the standard one for MSigDB signatures. As the other parameters, this object can be constructed at runtime using the dedicated upload widget.

All these modalities are equally valid for launching `ideal()`:

```
# all objects are precomputed and passed as parameters
ideal(dds_obj = dds, res_obj = res, annotation_obj = anno)
# the result object is constructed at runtime
ideal(dds_obj = dds)
# dds and res are computed at runtime, optionally with annotation - design is specified in the UI widget
ideal(countmatrix = countmatrix, expdesign = expdesign)
# the simplest form where all the required objects are assembled in the Shiny session - the UI guides the user sequentially to derive all the needed components
ideal()
```

These commands apply if the user launches `ideal` locally.
Other modalities of using `ideal` are described in a specific section of the main vignette (https://bioconductor.org/packages/release/bioc/vignettes/ideal/inst/doc/ideal-usersguide.html#3\_using\_the\_application).

#### 2.2 The demo dataset for this document

In this Section, we will be retrieving the experimental data (metadata) for the `macrophage` dataset, mentioned in the Introduction Section 1.

Here, we load the `SummarizedExperiment` object for the gene-level expression matrix, derived after summarizing via `tximport` the transcript level expression quantifications, obtained by running Salmon on the samples.

```
data("gse")
gse_macrophage <- gse
```

In the next step we generate `dds`, i.e. a `DESeqDataSet` object, which can be considered as the main input format to `ideal()`.
We specify the `design` parameter to be equal to `line + condition`, with `condition` specified last, as this is the one selected by default by *DESeq2* in its framework to generate the results.
This will allow us to compute the differences in expression levels with respect to the `condition`, while accounting for the cell `line` of origin.

```
dds_macrophage <- DESeqDataSet(se = gse_macrophage, 
                               design = ~line + condition)
dds_macrophage
# class: DESeqDataSet 
# dim: 58294 24 
# metadata(7): tximetaInfo quantInfo ... txdbInfo version
# assays(3): counts abundance avgTxLength
# rownames(58294): ENSG00000000003.14 ENSG00000000005.5 ...
#   ENSG00000285993.1 ENSG00000285994.1
# rowData names(2): gene_id SYMBOL
# colnames(24): SAMEA103885102 SAMEA103885347 ... SAMEA103885308
#   SAMEA103884949
# colData names(15): names sample_id ... condition line
```

Alternatively, one could provide the combination of count matrix and experimental metadata, supplied either as objects, or as text files at runtime.

Either way, we first work on the `rowData` slot of the `dds_macrophage`.

- We transform the original IDs from Gencode to Ensembl (we do this for better interoperability later on with the annotation packages), adding an extra column, `ensembl_id`
- We replace the original Gencode IDs with the Ensembl IDs in the `rownames`

```
rowData(dds_macrophage)$ensembl_id <- gsub('\\..+$', '', rownames(dds_macrophage))
# alternatively, one can use (for human, where the relevant part of the ID is composed by the first 15 characters)
# rowData(dds_macrophage)$ensembl_id <- substr(rownames(dds_macrophage),1,15)
rowData(dds_macrophage)
# DataFrame with 58294 rows and 3 columns
#                               gene_id      SYMBOL      ensembl_id
#                           <character> <character>     <character>
# ENSG00000000003.14 ENSG00000000003.14      TSPAN6 ENSG00000000003
# ENSG00000000005.5   ENSG00000000005.5        TNMD ENSG00000000005
# ENSG00000000419.12 ENSG00000000419.12        DPM1 ENSG00000000419
# ENSG00000000457.13 ENSG00000000457.13       SCYL3 ENSG00000000457
# ENSG00000000460.16 ENSG00000000460.16    C1orf112 ENSG00000000460
# ...                               ...         ...             ...
# ENSG00000285990.1   ENSG00000285990.1          NA ENSG00000285990
# ENSG00000285991.1   ENSG00000285991.1      RAET1E ENSG00000285991
# ENSG00000285992.1   ENSG00000285992.1          NA ENSG00000285992
# ENSG00000285993.1   ENSG00000285993.1          NA ENSG00000285993
# ENSG00000285994.1   ENSG00000285994.1          NA ENSG00000285994
rownames(dds_macrophage) <- rowData(dds_macrophage)$ensembl_id
```

Once this has been completed, you could start `ideal()` like this:

```
ideal(dds_macrophage)
```

For the sake of demonstrating an alternative entry point for `ideal()`, we export the counts matrix (derived from the estimated counts via Salmon) and the metadata to a text format.

```
write.table(counts(dds_macrophage),file = "idealinput_counts_macrophage.txt",
            quote = FALSE, col.names = colnames(dds_macrophage), sep = ",")
write.table(colData(dds_macrophage),file = "idealinput_metadata_macrophage.txt",
            quote = FALSE, col.names = colnames(colData(dds_macrophage)), sep = ",")
```

**Important:** The `macrophage` dataset described here differs from the one which is embedded in the demo instance of the app.
For simple reasons of size and startup time, *ideal* comes with the `airway` dataset, which has been generated in the work of (Himes et al. 2014), and is provided in the Bioconductor *airway* package as pre-computed matrix of raw counts.
This format is more efficient for the purpose of a smooth start, and the `airway` dataset is also the one that the standard tours refer to for guiding the user through the different offered features.

This document does refer to a more recent dataset, `macrophage`, processed according to the guidelines proposed by (Soneson, Love, and Robinson 2016).

### 3 Using `ideal()` - from uploading the count matrix all the way to downstream analyses

A distinctive trait of *ideal* is the possibility to perform most of the required steps, also described in the next Section (Section 4), with the ease of interactivity *yet* retaining the possibility to generate a fully-fledged R Markdown report, which gets knitted accessing the latest status of the reactive values when using the app.

This Section provides a step by step guide, with a number of screenshots, designed to guide the user through an exemplary full session, on the gene-level quantifications (counts, derived from transcript abundances) for the `macrophage` dataset.

If you followed the previous Section (Section 2.2) of this document, you will have available in your working directory all the ingredients to run a complete analysis with ideal, starting from the initial call with…

```
library("ideal")
ideal()
```

In the **Welcome** tab you can start familiarizing with the user interface, where the functionality is separated in the different tabs.
On the left you’ll see the sidebar, where the control widgets that affect multiple features are placed.
You can also spot out the Quick Viewer, just above the button to start the first introduction tour - we encourage all first users to click on it and follow through the steps (using the keyboard arrows is enabled).

Each tab has an additional dedicated collapsible help element (with a textual description) and a specific tour to showcase the features, with a “learning-by-doing” approach, which can be most efficient in complex web applications.

#### 3.1 Data Setup & Overview

Start uploading your count matrix and the experimental design matrix, using the file upload widgets inside the red box (Step 1, see Fig. 3.1).
You will use the two text files, `idealinput_counts_macrophage.txt` and `idealinput_metadata_macrophage.txt`, which you generated in the steps above (2.2).

The `macrophage` dataset is composed of 24 different samples, where cells from different lines and conditions have been sequenced in the scope of the original project.

If you did this correctly, this will populate the two collapsible boxes below (the Quick Viewer on the left sidebar will accordingly show two green checks), and also display a new box (Step 2), colored in yellow (Fig. 3.2).
Information on the format requirements of these text files is provided in the modal triggered by the question mark button.

Figure 3.1: Screenshot of the Welcome tab. Here, Step 1 is displayed, where both text files for the counts and the experimental metadata have been uploaded. On the lower left side, the collapsible element displays a portion of the uploaded expression matrix.

After this, you’ll have to specify the design and thus create the `DESeqDataSet` object (Fig. 3.2).
This is at least one of the column names of the provided experimental metadata information.
Select `line` and `condition` and click on the green action button (“Generate the dds object”).

Figure 3.2: Screenshot of the Welcome tab, after creating the DESeqDataset object in Step 2. The design ‘line + condition’ has been specified in the dedicated input widget.

In the `macrophage` dataset, we have 6 levels for the cell line, and 4 different experimental conditions.
In this document, we will focus on the `IFNg` vs `naive` comparison, where cells were either treated with Interferon-gamma, or left untreated.

Your selection will reflect in the formula displayed as `~line + condition`: this will enable you for example to estimate the effect size of the `condition` variable, while controlling for the cell `line` (Fig. 3.3).

Figure 3.3: Screenshot of the Optional steps in the Welcome tab, where the user just created the annotation object with the org.Hs.eg.db package.

Once you generated the `DESeqDataSet` object, you will see the ValueBox on top turn green, and also the sidebar field for it will get a green check mark.
You will be prompted by two optional steps (lower half of Fig. 3.2), where you can:

- create an Annotation object: this is especially useful if you are providing ENSEMBL or Gencode identifiers, which are stable over time but not human-readable such as HGNC gene symbols. Although optional, this step is highly recommended, since some functionality of *ideal* requires this object.
- remove samples from the dataset at hand (if you need to subset it, or if some outlier have been identified after a proper exploratory data analysis session, which you can e.g. perform also with the *pcaExplorer* package).

To retrieve the annotation for the `macrophage` dataset, simply select “Human” from the dropdown and “ENSEMBL” as id type.
This will load the required annotation package (*org.Hs.eg.db*), and pressing the action button will actually create the desired object - again, the corresponding ValueBox will turn green, and the Quick Viewer will get a green check icon.

Figure 3.4: Screenshot of Step 3 in the Welcome tab, where it is possible to run the main DESeq2 function - estimating size factors, estimating dispersions, and performing the testing. Once completed, a diagnostic plot can be displayed in the collapsible element.

At this point, you are all set to run the main function of the `DESeq2` framework, and you do so in Step 3 (green box), with the option of parallelizing the operations if multiple cores are available (Fig. 3.4).
Once this is completed, you can inspect the mean-dispersion plot as a diagnostic check by expanding the collapsible element below.

If you accidentally navigate to tabs where their content still needs to be generated, you will see the output of a conditional panel, which will point you towards the missing objects (Fig. 3.5).

Once the `dds` and the annotation have been specified, click on the “Run DESeq!” button.

Figure 3.5: Screenshot of the Extract Results tab, as it would be shown if the underlying required objects have not yet been computed.

From now on, you can continue in the **Counts Overview** tab (Fig. 3.6).
This features an interactive table which can display raw, normalized, and log-normalized values for all the genes and samples in the data.

A summary for the expressed features is reported below, and you can use a threshold for either criterium to filter out the lowly expressed genes (which can also reduce the computation time without impacting the quality of the results, if you e.g. completely exclude genes that are not detected in any sample).

For the `macrophage` dataset, in the “Basic summary for the counts” filter out genes that have not been detected in any sample.

Figure 3.6: Screenshot of the Counts Overview tab, where a table of the raw counts is displayed. Notice how two of the value boxes are shown in green, as the underlying objects have already been generated - only the DE data frame still needs to be computed.

A sample-to-sample scatterplot matrix (Fig. 3.7) can display an overview on the similarity across all individual samples without losing the information on the single features - this can be quite useful for detecting unexpected patterns for subset of genes (Rutter et al. 2019).

Figure 3.7: Screenshot of the scatterplot matrix for the 24 samples of the macrophage dataset, where a subset of 1000 genes has been used to reduce the time for the creation of the plot.

#### 3.2 Extracting results and visualizing them

If you got familiar with the `ideal()` app, you should recognize the collapsible element with some help and the button to start an introJS-based tour also in the **Extract Results** tab (Fig. 3.8).

First and foremost: you should select the alpha level for significance, which will define the threshold for calling a gene differentially expressed (depending on the specified null hypothesis).
Common values are 0.05 (default), 0.1 (more liberal), or 0.01 (more stringent).

For the `macrophage` set, we are interested in the IFNg vs naive comparison, so we select `condition` as experimental factor in the dropdown, and then select `IFNg` (numerator) versus `naive` (denominator) in the widgets below it.

You can leave the value for the False Discovery Rate set to default of 0.05.

If your selection is valid, a green “Extract the results” button will show up, and clicking on that will generate the full DE results table - again, its valueBox will become green, and the Quick Viewer will show all green checkmarks (Fig. 3.9).

Figure 3.8: Screenshot of the Extract Results tab, where the user is contrasting the IFNg treated cells vs naive cells, for the condition experimental variable. All other widgets have been set to the default values. Since the macrophage condition variable has more than two levels, the right side is displaying the possibility to perform an ANOVA-like analysis (e.g. contrasting ‘line + condition’ vs ‘line’).

In the `macrophage` dataset, more than 6000 genes have been detected as differentially expressed for the IfnG treatment vs naive contrast.
*IRF1*, *IL18BP*, and *GBP2* are for example the top regulated genes - sorted by adjusted p-value.

If the experimental factor of interest has more than 2 levels (as `condition` in this case), you could also conduct an ANOVA-like test (on the right side) - for this, we kindly refer to the `DESeq2` main vignette (http://www.bioconductor.org/packages/release/bioc/vignettes/DESeq2/inst/doc/DESeq2.html).

Figure 3.9: Screenshot of the result summary, both as an overview and as an interactive DT table, with identifiers displayed as buttons that link to relevant databases.

The table that appears below the buttons is again interactive, and is by default sorted by the adjusted p-value (Fig. 3.9).
If one provides the annotation object as recommended, the Ensembl identifiers and gene symbols become clickable buttons that directly link to external databases for that feature, either ensembl.org or the NCBI Gene DB - this is a seamless way to retrieve additional info e.g. about location, relevant literature, or known associated diseases.
A series of diagnostic plots follows (Fig. 3.10), and include:

- a raw p-value histogram, useful for checking the assumption of uniform distribution under the null hypothesis, also stratified by mean expression value (relevant if one is using the Independent Hypothesis Weighting for adjusting the p-value)
- a histogram of the log2 fold change values, to show its distribution and identify anomalies such as highly skewed tails
- a Schweder-Spjøtvoll plot (Schweder and Spjøtvoll 1982), showing the ranked p-values: this is a graphical method to illustrate the Benjamini-Hochberg multiple testing adjustment procedure, with the intersection point defining the subset of genes for which the False Discovery Rate (FDR) is controlled at the chosen level.

Figure 3.10: Screenshot of the four different diagnostic plots for the result section.

In the **Summary Plots** tab, users can interact with the MA plot (log2FoldChange vs mean expression values) by brushing on it - this will trigger a zoomed version of it to be displayed, along with the gene symbols defined in the annotation object (Fig. 3.11).
Drilling one step deeper, one can click close to any gene in the zoomed section, and obtain below a plot of the expression values, split by the experimental factor of interest (or any combination thereof).
That same gene is also searched in the Entrez database to display its full name and short description of it.

In the `macrophage` dataset, select some of the upregulated genes in the upper portion of the MA-plot.
The names for the selected genes will be highlighted in the focused panel as labels.
Clicking for example on the point for the *CXCL11* gene will display a boxplot for the expression values in all conditions, and additional info retrieved from the Entrez database.

Figure 3.11: Screenshot of the Summary Plots tab, where the MA-plot and its connected functionality is displayed. Brushing on the left plot, a zoomed version of it is displayed on the right, and the user can further narrow the focus on single features, shown in the UI elements below (a plot, and an information box).

Alternatively to the MA plot, users can also use a volcano plot (where the significance is directly plotted against the effect size, Fig. 3.12).

Figure 3.12: Screenshot of the volcano plot and the heatmaps in the Summary Plots section.

The subset of genes included in the rectangular selection is also displayed as heatmaps (both static and dynamic), and the underlying data is contained in the collapsible element at the bottom of Fig. 3.12.

If a subset of genes is known to be of interest, they can be explored further in the **Gene Finder** tab, where up to four can be shown singularly, or any amount can be annotated onto the MA plot (Fig. 3.13).
To avoid selecting many genes by hand (from the selectize widget in the sidebar), one can also upload the list as a plain text file (one column, one feature per row).

Figure 3.13: Screenshot of the Gene Finder tab, where three genes have been selected. The plot for CXCL11 is shown, together with the annotated MA-plot.

In the `macrophage` dataset, *CCL5*, *IFNGR1*, and *CXCL11* have been selected.
The boxplot for *CXCL11* is displayed in the third panel, above the annotated MA-plot, where all features are labelled.

#### 3.3 All the way downstream: functional analysis and exploring gene signatures

The **Functional Analysis** tab (Fig. 3.14) is a one-stop-shop to perform overrepresentation analysis (ORA) with a variety of methods:
- simple ORA as in *limma*, implemented in the `goana()` function
- correcting for gene length bias with *goseq*, as implemented in `ideal::goseqTable()`
- decorrelating the Gene Ontology graph structure with *topGO*), as in `pcaExplorer::topGOtable()`

These operations can be performed on the DE genes, either taken altogether (recommended, as most gene sets operate with coordinated changes in both up- and down-regulation), or split by the direction of expression change.
Additionally, up to two custom gene lists can be uploaded.
For simplifying the underlying operations, the genes are provided as symbols - therefore the importance to utilize the annotation object.

Figure 3.14: Screenshot of the Functional Analysis tab, where the analysis on all detected DE genes is performed for the ontology Biological Process via topGO. The table in the lower part is interactive, and has buttons to link to the external AmiGO database. By clicking on any row, the signature heatmap is generated.

The interactive table results provide external links to the AmiGO database via automatically generated buttons, and by clicking on one row of the tables generated via `topGO` (because of its ability to return non-redundant gene sets), users can display a heatmap where all the associated genes are displayed at once.

In the `macrophage` dataset, we take both up- and down-regulated genes, and look for enriched functions in the Biological Process ontology.
Click on the button to perform the analysis with `topGO`, and wait for the operation to be completed.
Unsurprisingly, we see the “interferon-gamma-mediated signaling pathway” term (\_GO:0060333\_) shows up among the most affected.
Clicking on it in the table displays the signature heatmap for the DE genes associated to it.

When more custom gene lists are uploaded, it is easy to represent their overlap with a Venn Diagram of selected subsets.

Figure 3.15: Screenshot of the Signature Explorer tab, after having completed the preprocessing required steps (uploading gmt file, computing variance stabilized data, matching identifiers, selecting signature, and controlling the plot output object).

The **Signature Explorer** (Fig. 3.15) extends the exploration of signature databases by accepting any text file adherent to the `gmt` (gene matrix transposed) format, which is commonly distributed e.g. by the widely used MSigDB database (http://software.broadinstitute.org/gsea/msigdb/index.jsp), or the WikiPathways database (http://data.wikipathways.org/). In this example, we will use the human hallmark gene sets (`h.all.v7.0.symbols.gmt`) and the c5 collection for Gene Ontology Biological Processes (`c5.bp.v7.0.symbols.gmt`) - you can retrieve them from http://software.broadinstitute.org/gsea/downloads.jsp (requires a login).

The first step in this case is to upload the `gmt` file, and the value box will report the number of signatures contained; directly after that, you can compute once the variance stabilized transformed data, which is amenable to visualization because of its homoskedastic behavior.

Figure 3.16: Screenshot for another MSigDB signature on the macrophage dataset, ‘Antigen processing and presentation of exogenous peptide antigen via MHC Class I’ from the c5.bp collection, version 7.0.

After this, it is important to match the identifier types - select “ENSEMBL” for your `dds` data, and “SYMBOL” for the signatures (you can recognize it from the filename, but it is always good practice to inspect the text file in an editor), with the `org.Hs.eg.db` package building the foundation for the conversion.
Next, select from the dropdown your signature of interest (you can exploit the autocompletion feature), and opt whether to display all its members, or just the ones detected as DE in your analysis; the lower left well panel contains some widgets for controlling the final aspect of the heatmap (with mean centering strongly recommended for better identifying the patterns of expressions across samples, Fig. 3.16).

Figure 3.17: Screenshot of the Signature Explorer tab, while the user is taking the guided introJS-based tour of the web application for the dedicated functionality.

In the `macrophage` dataset, the “GO\_ADAPTIVE\_IMMUNE\_RESPONSE” (defined in the `c5.bp.v7.0.symbols.gmt` file) is among the terms detected as overrepresented.
The heatmap for its components can be seen in Fig. 3.15.

This is probably best done by following the tour, as shown in Fig. 3.17.

#### 3.4 Exporting your analyses

Most of the output content generated in `ideal()` can be exported with a click on a download button, and either a plot (in quality-ready vectorial format) or a table (in plain text) can be generated.

Still, where `ideal()` excels in supporting reproducible research happens in the **Report Editor** tab (Fig. 3.18).
A template report in RMarkdown is part of the package itself, and constitutes the backbone upon which `ideal()` renders a fully fledged HTML report, previewed in the app itself, and can also be downloaded for storage or sharing among collaborators (Fig. 3.19 and Fig. 3.20).

Figure 3.18: Screenshot of the Report Editor tab, where the content of the template report is shown in the editor, and is being rendered - notice in the lower right corner the notification on the progress of the compilation.

Notably, experienced users can replace this template, or even modify and extend live during runtime, via the AceEditor in the “Edit Report” subtab (Fig. 3.18).
Options for the output document and the editor are in the collapsible elements.

In case users need visualizations not included in `ideal()`, the `dds` and `res` objects can be combined together (with the `res` becoming a nested `colData` component), and then exported to a single serialized `.rds` file (“Export as serialized SummarizedExperiment”, on the right side of Fig. 3.18), which can afterwards be seamlessly fed to the *iSEE* package, which also supports the combination of interactivity and reproducibility - see more on this at the demo instance of `iSEE` available at http://shiny.imbei.uni-mainz.de:3838/iSEE/.

Figure 3.19: Screenshot of the rendered report, as it is displayed in the tabbed content for a preview. In this image, the focus is on the initially provided parameters and objects.

Figure 3.20: Screenshot of the rendered report, focused on the intermediate output, as specified in the Gene Finder tab.

Additionally, the entire state of the app and its reactive objects can be exported to binary `.RData` objects, by clicking on the button accessed from the cogs icon (in the header of the app).
Exiting `ideal()` also saves these objects as environments, which can be accessed later in the prosecution of the offline analysis.
These buttons are accessed by clicking on the cog icon, shown e.g. in Fig. 3.18 above the green value boxes.

This concludes the analysis steps via the `ideal()` app.
Of course, users can navigate back to the previous tabs to investigate more in detail particular aspects, or iterate at will particular exploratory operations.

### 4 The main analysis - without `ideal()`

Experienced users might be familiar with the set of commands referring to the standard RNA-seq workflow, beautifully exemplified e.g. in https://master.bioconductor.org/packages/rnaseqGene/ or in https://bioconductor.org/packages/RNAseq123/ (Love et al. 2015; Law et al. 2016).

We can start doing some exploratory analysis and visualization on the expression matrix, after normalization and appropriate data transformations.
For this purpose, another package which might come in very handy is *pcaExplorer* (Marini and Binder 2019), which can be considered the counterpart of *ideal* for these preliminary steps, with a focus on Principal Components.

```
# prefiltering the dataset
dds_macrophage
nrow(dds_macrophage)
dds_macrophage <- dds_macrophage[ rowSums(counts(dds_macrophage)) > 0, ] # detected in at least one sample
nrow(dds_macrophage)

# normalizing
dds_macrophage <- estimateSizeFactors(dds_macrophage)
# transforming, and subsequently performing EDA
vsd <- vst(dds_macrophage)
rld <- rlog(dds_macrophage)

pcaExplorer::pcaplot(vsd, intgroup = "condition")
pcaExplorer::pcaplot(vsd, intgroup = "line", ellipse = FALSE)

sampleDistMatrix <- as.matrix( dist(t(assay(vsd))) )
colnames(sampleDistMatrix) <- rownames(sampleDistMatrix) <- 
  paste0(vsd$condition, "_", vsd$line)
pheatmap(sampleDistMatrix)
```

The `DESeq2` pipeline can be run with the following lines, and some summary info can be extracted.

```
# running the DE pipeline
dds_macrophage <- DESeq(dds_macrophage)

# comparing e.g. interferon gamma treated samples VS naive ones, with a strict log fold change threshold
res_macrophage <- results(dds_macrophage, contrast=c("condition","IFNg","naive"),
               lfcThreshold=1, alpha=0.01)
summary(res_macrophage)
# adding the gene symbols back to the result object
res_macrophage$gene_name <- rowData(dds_macrophage)$SYMBOL

DESeq2::plotMA(res_macrophage, ylim=c(-10,10))
```

We would then be proceeding with some functional enrichment analysis.

```
resOrdered <- as.data.frame(res_macrophage[order(res_macrophage$padj),])
de_df <- resOrdered[resOrdered$padj < 0.01 & !is.na(resOrdered$padj),]
de_symbols <- de_df$gene_name
bg_symbols <- rowData(dds_macrophage)$SYMBOL

# with topGO, from pcaExplorer
topgoDE_macro <- 
  pcaExplorer::topGOtable(DEgenes = de_symbols, 
                          BGgenes = bg_symbols,
                          ontology = "BP",
                          mapping = "org.Hs.eg.db",
                          geneID = "symbol",
                          addGeneToTerms = TRUE)
DT::datatable(topgoDE_macro)

# with goseq, from ideal
goseqDE_macro <- ideal::goseqTable(
  de.genes = rownames(de_df),
  assayed.genes = rownames(dds_macrophage),
  testCats = "GO:BP")
DT::datatable(goseqDE_macro)

# with clusterProfiler
ego_macro <- enrichGO(gene = de_symbols,
                      universe = bg_symbols,
                      OrgDb = org.Hs.eg.db,
                      keyType = "SYMBOL",
                      ont = "BP",
                      pAdjustMethod = "BH",
                      pvalueCutoff = 0.01,
                      qvalueCutoff = 0.05)
head(ego_macro)
emapplot(ego_macro)
```

For generating signature heatmaps of the gene set of interest, some bespoke lines of code might be necessary.
Users can explore the template report and source code of `ideal` to obtain similarly fashioned graphics.

### Session info

```
sessionInfo()
# R version 3.6.0 (2019-04-26)
# Platform: x86_64-apple-darwin15.6.0 (64-bit)
# Running under: macOS Sierra 10.12.6
# 
# Matrix products: default
# BLAS:   /Library/Frameworks/R.framework/Versions/3.6/Resources/lib/libRblas.0.dylib
# LAPACK: /Library/Frameworks/R.framework/Versions/3.6/Resources/lib/libRlapack.dylib
# 
# locale:
# [1] en_US.UTF-8/en_US.UTF-8/en_US.UTF-8/C/en_US.UTF-8/en_US.UTF-8
# 
# attached base packages:
# [1] parallel  stats4    stats     graphics  grDevices utils     datasets 
# [8] methods   base     
# 
# other attached packages:
#  [1] ideal_1.11.0                topGO_2.38.1               
#  [3] SparseM_1.78                GO.db_3.10.0               
#  [5] graph_1.64.0                pcaExplorer_2.13.0         
#  [7] bigmemory_4.5.36            knitr_1.26                 
#  [9] clusterProfiler_3.14.2      pheatmap_1.0.12            
# [11] org.Hs.eg.db_3.10.0         AnnotationDbi_1.48.0       
# [13] DESeq2_1.26.0               tximeta_1.4.2              
# [15] SummarizedExperiment_1.16.1 DelayedArray_0.12.1        
# [17] BiocParallel_1.20.1         matrixStats_0.55.0         
# [19] Biobase_2.46.0              GenomicRanges_1.38.0       
# [21] GenomeInfoDb_1.22.0         IRanges_2.20.1             
# [23] S4Vectors_0.24.1            BiocGenerics_0.32.0        
# [25] magrittr_1.5                macrophage_1.3.1           
# 
# loaded via a namespace (and not attached):
#   [1] rappdirs_0.3.1           rtracklayer_1.46.0       AnnotationForge_1.28.0  
#   [4] pkgmaker_0.27            tidyr_1.0.0              ggplot2_3.2.1           
#   [7] acepack_1.4.1            bit64_0.9-7              data.table_1.12.8       
#  [10] rpart_4.1-15             RCurl_1.95-4.12          AnnotationFilter_1.10.0 
#  [13] doParallel_1.0.15        GenomicFeatures_1.38.0   cowplot_1.0.0           
#  [16] RSQLite_2.1.5            europepmc_0.3            bit_1.1-14              
#  [19] enrichplot_1.6.1         BiocStyle_2.14.2         xml2_1.2.2              
#  [22] httpuv_1.5.2             assertthat_0.2.1         viridis_0.5.1           
#  [25] xfun_0.11                tximport_1.14.0          hms_0.5.2               
#  [28] evaluate_0.14            promises_1.1.0           IHW_1.14.0              
#  [31] progress_1.2.2           caTools_1.17.1.3         dbplyr_1.4.2            
#  [34] Rgraphviz_2.30.0         igraph_1.2.4.2           DBI_1.1.0               
#  [37] geneplotter_1.64.0       htmlwidgets_1.5.1        purrr_0.3.3             
#  [40] crosstalk_1.0.0          dplyr_0.8.3              backports_1.1.5         
#  [43] bookdown_0.16            annotate_1.64.0          gridBase_0.4-7          
#  [46] biomaRt_2.42.0           vctrs_0.2.1              ensembldb_2.10.2        
#  [49] withr_2.1.2              ggforce_0.3.1            triebeard_0.3.0         
#  [52] checkmate_1.9.4          GenomicAlignments_1.22.1 fdrtool_1.2.15          
#  [55] prettyunits_1.0.2        cluster_2.1.0            DOSE_3.12.0             
#  [58] lazyeval_0.2.2           crayon_1.3.4             genefilter_1.68.0       
#  [61] pkgconfig_2.0.3          slam_0.1-47              tweenr_1.0.1            
#  [64] nlme_3.1-143             ProtGenerics_1.18.0      nnet_7.3-12             
#  [67] rlang_0.4.2              lifecycle_0.1.0          registry_0.5-1          
#  [70] bigmemory.sri_0.1.3      BiocFileCache_1.10.2     GOstats_2.52.0          
#  [73] polyclip_1.10-0          rngtools_1.4             Matrix_1.2-18           
#  [76] urltools_1.7.3           lpsymphony_1.14.0        base64enc_0.1-3         
#  [79] geneLenDataBase_1.22.0   ggridges_0.5.1           png_0.1-7               
#  [82] viridisLite_0.3.0        bitops_1.0-6             shinydashboard_0.7.1    
#  [85] KernSmooth_2.23-16       Biostrings_2.54.0        blob_1.2.0              
#  [88] rintrojs_0.2.2           stringr_1.4.0            qvalue_2.18.0           
#  [91] jpeg_0.1-8.1             gridGraphics_0.4-1       shinyAce_0.4.1          
#  [94] scales_1.1.0             memoise_1.1.0            GSEABase_1.48.0         
#  [97] plyr_1.8.5               gplots_3.0.1.1           bibtex_0.4.2.1          
# [100] gdata_2.18.0             zlibbioc_1.32.0          threejs_0.3.1           
# [103] compiler_3.6.0           RColorBrewer_1.1-2       d3heatmap_0.6.1.2       
# [106] Rsamtools_2.2.1          XVector_0.26.0           Category_2.52.1         
# [109] htmlTable_1.13.3         Formula_1.2-3            MASS_7.3-51.5           
# [112] mgcv_1.8-31              tidyselect_0.2.5         stringi_1.4.3           
# [115] shinyBS_0.61             highr_0.8                yaml_2.2.0              
# [118] GOSemSim_2.12.0          askpass_1.1              locfit_1.5-9.1          
# [121] latticeExtra_0.6-29      ggrepel_0.8.1            grid_3.6.0              
# [124] fastmatch_1.1-0          tools_3.6.0              rstudioapi_0.10         
# [127] foreach_1.4.7            foreign_0.8-74           gridExtra_2.3           
# [130] farver_2.0.1             ggraph_2.0.0             digest_0.6.23           
# [133] rvcheck_0.1.7            BiocManager_1.30.10      shiny_1.4.0             
# [136] Rcpp_1.0.3               later_1.0.0              httr_1.4.1              
# [139] colorspace_1.4-1         XML_3.98-1.20            splines_3.6.0           
# [142] RBGL_1.62.1              graphlayouts_0.5.0       ggplotify_0.0.4         
# [145] xtable_1.8-4             jsonlite_1.6             tidygraph_1.1.2         
# [148] UpSetR_1.4.0             zeallot_0.1.0            R6_2.4.1                
# [151] Hmisc_4.3-0              pillar_1.4.3             htmltools_0.4.0         
# [154] mime_0.8                 NMF_0.21.0               glue_1.3.1              
# [157] fastmap_1.0.1            DT_0.11                  codetools_0.2-16        
# [160] fgsea_1.12.0             lattice_0.20-38          tibble_2.1.3            
# [163] rentrez_1.2.2            curl_4.3                 BiasedUrn_1.07          
# [166] gtools_3.8.1             openssl_1.4.1            survival_3.1-8          
# [169] limma_3.42.0             rmarkdown_2.0            munsell_0.5.0           
# [172] DO.db_2.9                GenomeInfoDbData_1.2.2   iterators_1.0.12        
# [175] goseq_1.38.0             reshape2_1.4.3           gtable_0.3.0
```

### References

Alasoo, Kaur, Julia Rodrigues, Subhankar Mukhopadhyay, Andrew J. Knights, Alice L. Mann, Kousik Kundu, Christine Hale, Gordon Dougan, and Daniel J. Gaffney. 2018. “Shared genetic effects on chromatin and gene expression indicate a role for enhancer priming in immune response.” *Nature Genetics* 50 (3). Springer US: 424–31. https://doi.org/10.1038/s41588-018-0046-7.

Himes, B. E., Jiang, X., Wagner, P., Hu, et al. 2014. “RNA-Seq Transcriptome Profiling Identifies CRISPLD2 as a Glucocorticoid Responsive Gene that Modulates Cytokine Function in Airway Smooth Muscle Cells.” *PLoS ONE* 9 (6): e99625. http://www.ncbi.nlm.nih.gov/pubmed/24926665.

Law, Charity W., Monther Alhamdoosh, Shian Su, Gordon K. Smyth, and Matthew E. Ritchie. 2016. “RNA-seq analysis is easy as 1-2-3 with limma, Glimma and edgeR.” *F1000Research* 5 (0): 1408. https://doi.org/10.12688/f1000research.9005.1.

Love, Michael I., Simon Anders, Vladislav Kim, and Wolfgang Huber. 2015. “RNA-Seq workflow: gene-level exploratory analysis and differential expression.” *F1000Research* 4: 1070. https://doi.org/10.12688/f1000research.7035.1.

Marini, Federico, and Harald Binder. 2019. “pcaExplorer: an R/Bioconductor package for interacting with RNA-seq principal components.” *BMC Bioinformatics* 20 (1): 331. https://doi.org/10.1186/s12859-019-2879-1.

Rutter, Lindsay, Adrienne N. Moran Lauter, Michelle A. Graham, and Dianne Cook. 2019. “Visualization methods for differential expression analysis.” *BMC Bioinformatics* 20 (1). BMC Bioinformatics: 1–31. https://doi.org/10.1186/s12859-019-2968-1.

Schweder, T, and E Spjøtvoll. 1982. “Plots of P-values to evaluate many tests simultaneously.” *Biometrika* 69 (3): 493–502. https://doi.org/10.1093/biomet/69.3.493.

Soneson, Charlotte, Michael I. Love, and Mark D. Robinson. 2016. “Differential analyses for RNA-seq: transcript-level estimates improve gene-level inferences.” *F1000Research* 4 (0): 1521. https://doi.org/10.12688/f1000research.7563.2.
